# Engineered nickel bioaccumulation in *Escherichia coli* by NikABCDE transporter and metallothionein overexpression

**DOI:** 10.1101/2022.08.26.505507

**Authors:** P. Diep, H. Shen, J. A. Wiesner, N. Mykytczuk, V. Papangelakis, A. F. Yakunin, R. Mahadevan

**Affiliations:** BioZone – Centre for Applied Bioscience and Bioengineering, Department of Chemical Engineering and Applied Chemistry, University of Toronto, Toronto, ON, Canada; MIRARCO Mining Innovation, Laurentian University, Sudbury ON, Canada; Centre for Environmental Biotechnology, School of Natural Sciences, University of Bangor, Wales, United Kingdom; Institute of Biomedical Engineering, University of Toronto, Toronto, ON, Canada

**Author notes:** Correspondence: Radhakrishnan Mahadevan.

**Keywords:** industrial biotechnology, wastewater sampling, genetic engineering, metal recovery, ABC transporter

## Abstract

Mine wastewater often contains dissolved metals at concentrations too low to be economically extracted by existing technologies, yet too high for environmental discharge. The most common treatment is chemical precipitation of the dissolved metals using limestone and subsequent disposal of the sludge in tailing impoundments. While it is a cost-effective solution to meet regulatory standards, it represents a lost opportunity. In this study, we engineered *Escherichia coli* to overexpress its native NikABCDE transporter and a heterologous metallothionein to capture nickel at concentrations in local effluent streams. We found the engineered strain had a 7-fold improvement in the bioaccumulation performance for nickel compared to controls, but also observed a drastic decrease in cell viability due to metabolic burden or inducer (IPTG) toxicity. Growth kinetic analysis revealed the IPTG concentrations used based on past studies lead to growth inhibition, thus delineating future avenues for optimization of the engineered strain and its growth conditions to perform in more complex environments.

## INTRODUCTION

Biomining, the use of biotechnology for mining applications has been gaining new traction with pilot-scale and commercial-scale demonstrations of microbial cultures performing bioleaching processes to aid in the recovery of lithium and other metals from low-grade ores and mine waste (1). One form of mine waste are liquid effluents bearing dissolved metals too low in concentration (≤ low ppm) to be economically extracted with traditional practices, but in excess of regulatory standards prohibiting environmental discharge without some form of treatment. Chemical precipitation of metals by sulfide or hydroxide addition is the primary form of treatment, yet it generates secondary sludge waste that must be disposed of in tailing impoundments. Biomining includes processes like biosorption (*i*.*e*., the adsorption of dissolved metals to the surface of cells) and bioprecipitation (*i*.*e*., altering the redox state of dissolved metals to decrease their solubility in water), which have been extensively reported in the literature as alternatives (Fig. 1A). While some researchers have focused on leveraging microbes’ natural capabilities for biomining, others have explored the expansion the microbes’ capabilities through the latest advances in synthetic biology (2–4).

**Figure 1.**
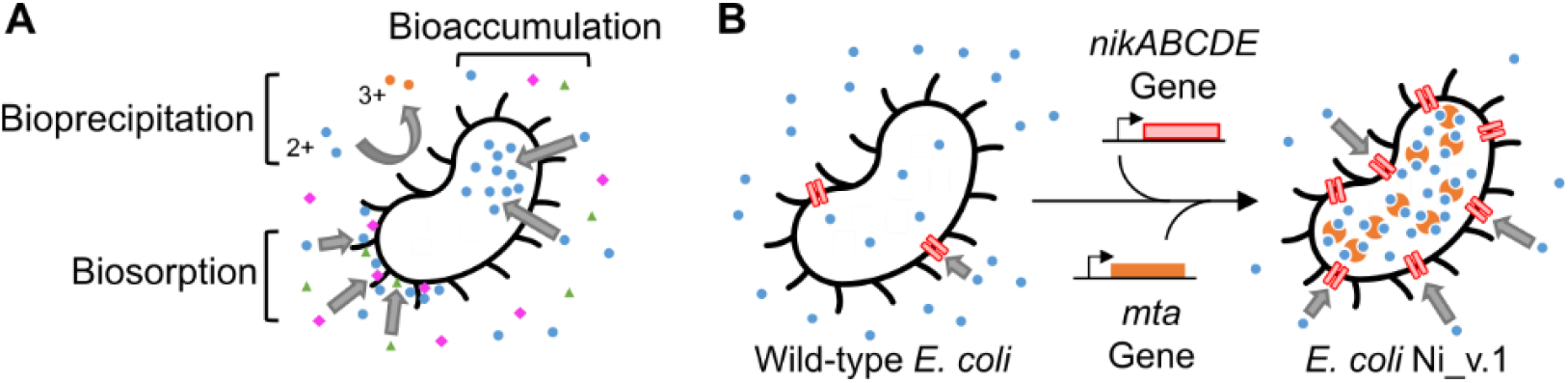
Ni_v.1 design. (A) Important cell-metal ion interactions for metal removal bioprocesses. Different colours shapes represent different metals. Biosorption involves metals adhering to moieties on the surface of the cell. Bioprecipitation involves a metal redox change to an insoluble form driven by extracellular and membrane-bound enzymes. Bioaccumulation involves the specific import of metals into the cytosol. (B) Genetic engineering design to enhance *Escherichia coli*’s bioaccumulation performance for importing (*nikABCDE*) and storing (*mta*) dissolved nickel (blue circles).

Synthetic biology is a growing discipline that seeks to design microorganisms and biomolecular systems by combining decades of life sciences research with design principles and tools from math, physics, and engineering. Biosorption and bioprecipitation have benefited from advances in synthetic biology. For example, the surface of microbes have been designed to bear lanthanide-binding peptides to enhance their ability to adsorb dissolved rare earth elements (5). Other microbes have been designed to produce H_2_S to precipitate dissolved metals as nanoparticles (6). While these advances are promising, biosorption processes need to overcome selectivity issues since the surface of the cell directly interfaces with the aqueous environment and is therefore more susceptible to a molecular form of fouling. Bioprecipitation processes produce metal products with redox states that are insoluble in water. An increasingly important objective in the field, beyond bioremediation, is to recover the metals in a market-ready form. This means one would need to reverse the redox change to re-integrate the metal into existing purification processes, which would pose additional operation costs. A desirable bioprocess that can compete with existing technologies must therefore possess robust selectivity for specific dissolved metals while being able to smoothly integrate with existing hydrometallurgical processes.

We thus focused our attention on bioaccumulation. Microbes can naturally bioaccumulate metals selectively through highly specific protein transport systems evolved to capture metals from metal-scarce environments like the human gut (7). We previously reviewed studies where microorganisms were genetically engineered to express heterologous transporters and metal-binding “storage” proteins for the purpose of enhancing the hosts’ abilities to bioaccumulate metal in the cytosol and noted the absence of studies that used ABC transporters for the purpose of metal removal from liquid effluents (11). Bioaccumulation is a unique strategy to develop a bioprocess. While it is generally slower than biosorption, the cell membrane acts as a selectivity barrier, and the presence of transporters allow specific metals to enter the cytosol without the need for a change in the redox state. A critical system is the ATP-binding cassette (ABC) transporter, a class of active transporters that convert the chemical energy from nucleotide triphosphate (NTP) hydrolysis into mechanical energy that drives the movement of chemicals and ions across the lipid membrane (8–10). ABC transporters are comprised of two components: the solute binding protein (SBP) found in the periplasm or the extracellular space via lipid tether to the membrane, and the transmembrane permease complex baring additional cytosolic NTP-binding domains. Taken together, we briefly proposed in our review the potential for ABC transporters to be an advantageous design choice to enhance a cell’s bioaccumulation performance, defined as the mg_metal_/g_dry cell weight_ (or mg_x_/g_dcw_ where ‘x’ denotes the metal).

Conducting design-build-test-learn (DBTL) cycles is a common engineering practice within synthetic biology to iteratively improve strain performance towards an objective. In this study, we focused on initializing the first DBTL cycle and began by taking inspiration from early work by Deng *et al*. (2003, 2005) to design Ni_v.1, an *Escherichia coli* strain with enhanced bioaccumulation performance for nickel using a nickel-specific ABC transporter complemented with a protein-based storage system as described in Fig.1B (12, 13). Specifically, we built the strain by cloning the open-reading frame of *nikABCDE* from *E. coli* BL21(DE3) genomic DNA, and *mta* synthesized commercially into the T7-controlled dual expression vector pCDF-Duet, then transformed this plasmid into an *E. coli* BL21(DE3) to heterologously overexpress the transport-storage system. NikABCDE is an experimentally validated nickel-specific transporter (14), and it’s periplasmic NikA is especially well-characterized (15–18). MTA is a cysteine-rich metallothionein from *Pisum sativum* previously used successfully for enhancing bioaccumulation of nickel, cobalt, and mercury (11).

To guide our testing, we sampled real Canadian liquid mine effluent from the Sudbury, Ontario community and performed a simple compositional analysis to determine the future working constraints (*i*.*e*., metal concentrations, salinity, and pH) that Ni_v.1 would need to perform under. Since this was the first DBTL cycle, we benchmarked Ni_v.1’s performance in unbuffered 10 ppm NiCl_2_ solution (matching the concentration found in the Copper Cliff sample) to understand its performance in simplified ideal conditions (Fig. 2). An initial test of Ni_v.1 compared to its control strains revealed it had the highest bioaccumulation performance for nickel that was comparable to previously reported values in the literature. Finally, we found that induction conditions could be further optimized by reducing the IPTG concentration in favour of better growth kinetics. Based on these findings, we proposed future avenues of research in subsequent DBTL cycles for improving Ni_v.1’s bioaccumulation performance for nickel.

**Figure 2.**
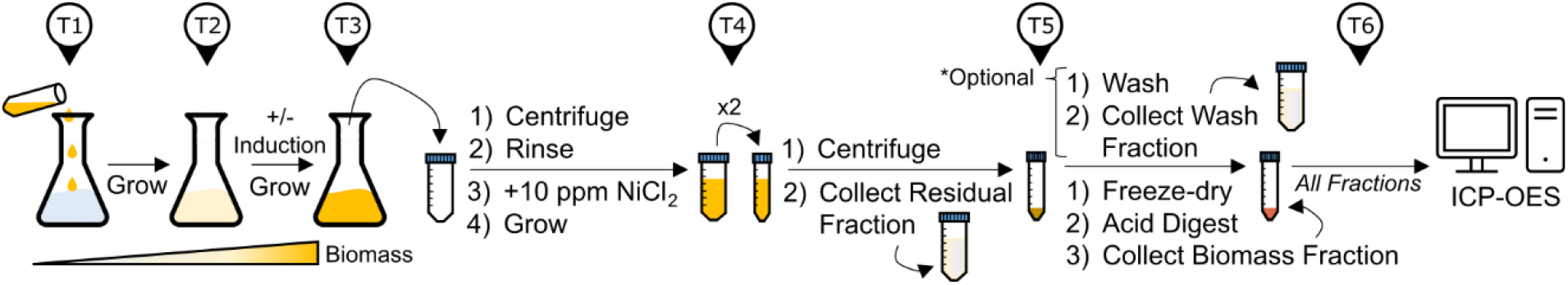
Shake flask workflow to assay for bioaccumulation performance. Timepoints denoted above (T1, T2, *etc*.) to indicate chronological order of steps. The culture begins at T1 inoculation and is grown until protein expression is ready to be induced at T2. After the induction period, at T3 the cell biomass is isolated by centrifugation and resuspended in the nickel solution for the cells to bioaccumulate the nickel. The optical density (O.D.) of the culture can be measured at T1→T3. At T4, the cells are collected again by centrifugation and the supernatant is collected as the Residual Fraction. At T5, the nickel-loaded cell pellet can be washed to collect a Wash Fraction that enables the differentiation of the nickel biosorped to the surface of the cell from the nickel bioaccumulate inside the cytosol. The cells are collected by centrifugation to obtain the Biomass Fraction and freeze-dried to obtain the dry cell weight. The dried cell pellets are acid digested overnight and analyzed by ICP-OES at T6.

## RESULTS AND DISCUSSION

### Test 0: Estimating Ni_v.1 working conditions based on aqueous streams from a mine

Sudbury is home to the century-old Vale Copper Cliff site, one of the world’s largest active integrated mining complexes. The site includes grinding and milling stations, smelters, refineries, and – critical to this study – tailings deposition sites for both solid and liquid waste. To maintain their water balance, liquid effluents are treated by lime addition to precipitate the metals and neutralize the aqueous stream prior to environmental discharge. Interestingly, these aqueous streams run through several Sudbury communities as part of the Junction Creek Watershed. We sought to ground the testing of Ni_v.1 in real-world constraints, so we identified two water sampling sites of interest: Nolin Creek (N.C.) in the northern part of Sudbury, and a Copper Cliff stream (C.C.) in the southern part (Fig. S1 for aerial view). N.C. was located ∼2.3 km from its origin in the Vale Copper Cliff mine and was downstream of a lime treatment plant (Fig. 3A). C.C. was located ∼1 km from its origin in the same mine, but was not treated yet since it was upstream of a lime treatment plant (Fig. 3C), thus explaining the acid mine drainage characteristics (*e*.*g*., reddish-orange iron precipitate and low pH).

**Figure 3.**
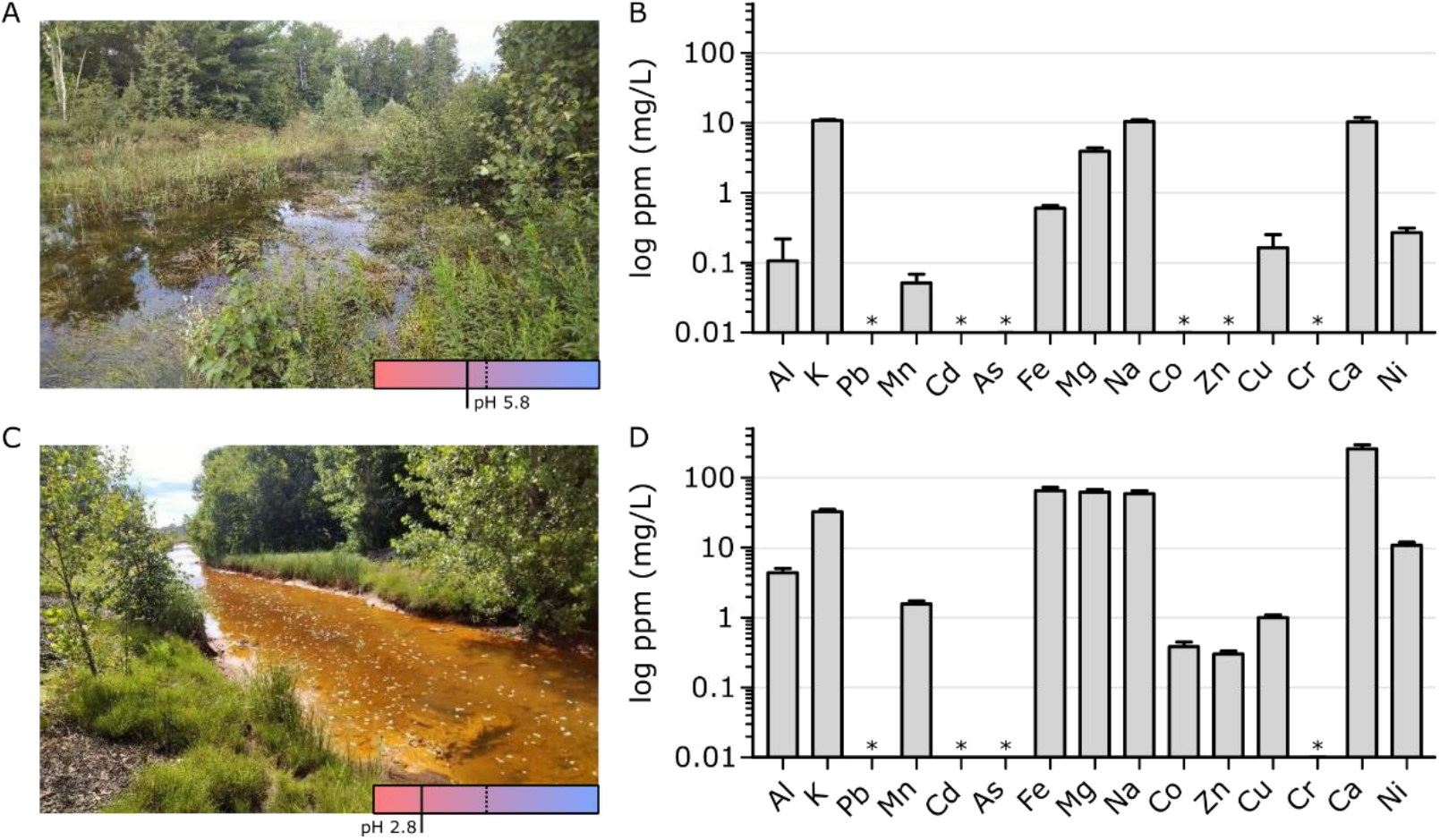
Metal analysis of aqueous streams originating from Vale Copper Cliff. (A) Westward view of sampling site at Nolin Creek (N.C.) and (B) its metal composition (n=3). Dotted line indicates pH 7 in the bottom right pH indicator box. C) Eastward view of an aqueous stream flowing from a Copper Cliff tailings dam (C.C) and (D) its metal composition (n=3). Asterisk (*) denotes measurements below the limit of detection. Metal measurements made using ICP-OES.

We obtained three samples at each of the two sites and analyzed their composition (Fig. 3). The C.C. samples (pH 2.84) were significantly more acidic than the N.C. samples (pH 5.83), thus leading to considerably distinct chemical conditions affecting the solubility of the metals (Table S1, Fig. 3B,D). Several metals (Al, Mn, Fe, Mg, Na, and Ca) were all at least one order of magnitude more concentrated in C.C. samples than N.C. samples (Fig. S2 for side-by-side comparison). Toxic metal ions (As, Pb, Cd, and Cr) were below the limit of detection at both sites, as were Co and Zn at N.C. Our element of interest nickel was 10.805 ± 1.118 ppm in the C.C. sample, which was a 40-fold higher than N.C. samples. Both sites were predominantly sulfate systems (>99%) with trace amounts of phosphates and no detectable chlorides, which reflected the sulfidic mineral geology native to Copper Cliff. The total positive charge concentrations in the samples from N.C. and C.C. were to 1.63 mM and 27.26 mM of e^-^, respectively. For the negative charge concentrations, the calculated values were 1.77 mM and 27.79 mM of e^-^. The small errors (2% and 8% for C.C. and N.C., respectively) in these charge balances indicate that most of the ions in the samples had been captured in this analysis.

Our goal in this first DBTL cycle was to confirm if Ni_v.1 had appreciable bioaccumulation performance for nickel compared to similarly engineered strains reported in literature. We previously found acidic or high salinity environments could reduce the nickel-binding affinity of a *Ec*NikA homologue called *Cc*SBPII from *Clostridium carboxidivorans*. Phosphate buffer, compared to more inert buffers like HEPES and MOPS, also reduced *Cc*SBPII’s nickel-binding affinity (19). Given the concentration of nickel in the C.C. samples and the above considerations, we prepared 10 ppm NiCl_2_ solutions to benchmark Ni_v.1’s bioaccumulation performance for nickel. We intentionally i) avoided adding salt (Na, K, Mg, Ca), ii) chose NiCl_2_ for a chloride anion system rather than phosphate or sulfate salts that would introduce polyatomic anions (PO_4_ ^3-^, SO_4_ ^2-^), iii) did not buffer the pH as that would require the addition of millimolar levels of buffering compounds that would not be found in nature, and iv) only included nickel in the simulated effluent since we were not studying bioaccumulation selectivity.

### Test 1: Benchmarking Ni_v.1 bioaccumulation performance for nickel

The workflow outlined in Fig.2 to measure bioaccumulation performance is laborious, so we performed flask experiments to determine whether Ni_v.1 would function as we expected before testing its reproducibility (Fig. S4). In the absence and presence of IPTG at 0.4 mM based on prior studies (12, 13), we compared the performance of Ni_v.1 with control strains that either had the MTA storage system (“MTA”), the NikABCDE transport system (NikABCDE), or neither system (*i*.*e*., “Vector Only”). Our observations suggested Ni_v.1 was indeed functional based on its bioaccumulation performance of 7.51 mg_Ni_/g_dcw_ in the presence of IPTG, which was 9.9-fold higher than its absence. In contrast, induction of the Vector Only strain did not lead to a notable change in the bioaccumulation performance, from 0.83 to 1.02 mg_Ni_/g_dcw_, which we expected. While the addition of IPTG also did not change the MTA control’s strains performance (from 0.31 to 0.20 mg_Ni_/g_dcw)_, it did increase the NikABCDE control’s performance by 3-fold from 1.08 to 3.31 mg_Ni_/g_dcw_. The 2.3-fold improvement in performance between the induced NikABCDE control and induced Ni_v.1 reflects the importance of having a storage system in the cytosol to sequester the accumulated metal and minimize loss through efflux pumps or cell lysis. Additionally, the 38-fold improvement between the induced MTA control and induced Ni_v.1 reflects the inability of our strain to use its native transport systems with the heterologous storage system to bioaccumulate nickel. Overall, Ni_v.1’s performance in this preliminary test is comparable to other engineered strains reported in the literature (*cf*. Test 2).

Throughout the workflow, we noticed that induced Ni_v.1 had less biomass at timepoint 3 (T3) than the non-induced Ni_v.1 control and the other controls. At T5, the appearance of the Biomass fraction for induced Ni_v.1 was also different from the other controls (Fig. S3). Instead of solid beige pellet, induced Ni_v.1 had a smaller, darker beige-black pellet with an amorphous pink layer of unknown material on top. We hypothesized the change in appearance was due to cell lysis resulting from a burden imposed by 10 ppm nickel or 0.4 mM IPTG. Next, we tested the reproducibility of Ni_v.1’s bioaccumulation performance using the Vector Only strain as the control and, considering these growth issues, we monitored the optical density (O.D.) of the cultures from T1 to T3.

### Test 2: Ni_v.1’s bioaccumulation performance and the associated burden is reproducible

We repeated the workflow for the Vector Only control and Ni_v.1 in the presence and absence of 0.4 mM IPTG (n=3, each). We also included an additional triplicate labelled “No Cells” where no inoculums were added at T1, but the samples were processed exactly the same as the test group. This was to check the sterility of nutrient media and to assess the nickel mass balance. No growth was observed in the No Cells control between T1-T3. The No Cell control experienced a 7.3 ± 2.1% loss of nickel, which we expected since there were several transferring and vortexing steps where solution adhering to pipettes and lids was not be collected (Fig. 4A). The test group experienced a 17.4 ± 1.7% loss of nickel, likely because any lost solution would also contain nickel-loaded cells (either biosorped to the surface or bioaccumulated into the cell) and thus amplify the losses. Nonetheless, the test group had relatively consistent losses of nickel between the samples, which enabled reliable cross-comparisons of their bioaccumulation performance.

**Figure 4.**
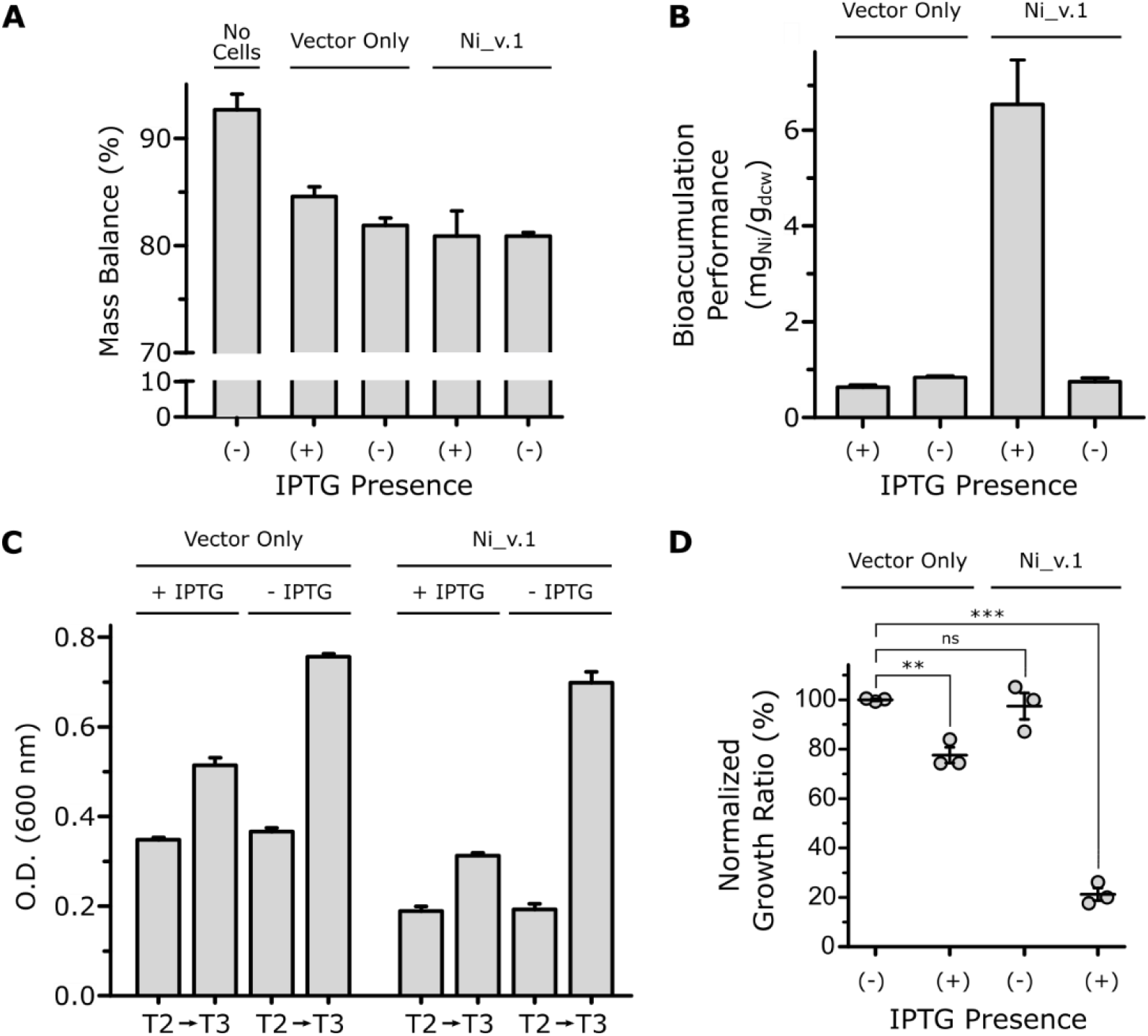
Ni_v.1 bioaccumulation performance and growth analysis. (A) Determination of mass balance across all fractions (*i*.*e*., residual, wash, and biomass) compared to a control with no cells initially added to the flasks intended for bioaccumulation cultures (n=3). (B) The bioaccumulation performance of wild-type *E. coli* compared to *E. coli* Ni_v.1 (n=3). (C) Comparison of growth rate between wild-type *E. coli* and Ni_v.1 before and after the timepoint where one half of the cultures for each strain are induced by the addition of IPTG (n=3). (D) Negative synergistic effect of IPTG induction and exposure to Ni^2+^ on Ni_v.1 (n=3).

The results of this test (Fig. 4B) were similar to observations in Fig.S4. Induced Ni_v.1 achieved a performance of 6.53 ± 1.47 mg_Ni_/g_dcw_, which was significantly higher (8.7-fold) than its non-induced control at 0.75 ± 0.11 mg_Ni_/g_dcw_ (unpaired two-tail t-test, p = 0.0051, t = 5.561). The induced and non-induced Vector Only control displayed no significant change: 0.63 ± 0.07 mg_Ni_/g_dcw_ and 0.92 ± 0.33 mg_Ni_/g_dcw_, respectively (unpaired two-tail t-test, p = 0.3009, t = 1.187). Compared to other enhanced nickel bioaccumulation strains reported in literature, and factoring in their starting concentration of nickel, Ni_v.1 has the second best performance (11). However, the best strain reported by Krishnaswamy & Wilson (2000) used a protocol that deployed a 10 mM phosphate buffer double-wash, compared to our more stringent 10 mM EDTA wash, which may have lead to an overestimation of their best strain’s performance (20). Nonetheless, standardization of protocols to measure bioaccumulation performance are needed since the contributions of biosorption can lead to overestimations of a bioaccumulation metric. The similarity of Ni_v.1’s performance to strains already reported in literature after one DBTL cycle demonstrates promising future iterations. We noted that Ni_v.1 was able to concentrate the nickel 175-fold with the potential to achieve >1000-fold concentration if the dried cell pellet was acid digested in <1 mL as opposed to the 5 mL that was used in this test.

The total nickel extraction by the cell from solution (Fig. S5), accounting for nickel in both the wash and biomass fractions, was approximately the same between the induced samples (60.9 ± 1.6%) and between the non-induced samples (46.6 ± 3.6%). However, when normalized to the mass of the dried cell weight (g_dcw_), induced Ni_v.1 had 6-to 9-fold higher nickel extraction from solution by bioaccumulation than the other controls. Together, these metrics indicated Ni_v.1 had the highest bioaccumulation performance, and the best removal of nickel from the 10 ppm NiCl_2_ solution specifically due to the bioaccumulation mechanism. However, Ni_v.1 again had weaker growth.

By following the growth kinetics of this test group, we quantified our prior observations in Test 1 (Table S2). Before resuspension in 10 ppm NiCl_2_ (Fig. 4C), we found the Ni_v.1 samples’ O.D. values at T2 were roughly half of the Vector Only controls’ O.D values (0.19 ± 0.09 and 0.36 ± 0.17, respectively). We attributed this inhibition to the burden imposed by the difference in the plasmid sizes since Ni_v.1 carries a larger 8.8 kbp plasmid for the production of the NikABCDE and MTA compared to the empty 3.8 kbp plasmid of the Vector Only control (21). From T2, we found the O.D. of the non-induced Vector Only controls and Ni_v.1 samples at T3 had increased approximately 207% (from O.D. 0.36 ± 0.01 to 0.76 ± 0.01) and 362% (from O.D. 0.19 ± 0.02 to 0.70 ± 0.03), respectively, which allowed Ni_v.1 to “catch-up” despite the initial lag at T2. In contrast, the induced groups only increased approximately 148% (from O.D. 0.35 ± 0.01 to 0.51 ± 0.02) and 165% (from O.D. 0.19 ± 0.01 to 0.31 ± 0.01), respectively, meaning the amount of induced Ni_v.1 biomass going into resuspension with 10 ppm NiCl_2_ for bioaccumulation was ∼60% less than least inhibited strain condition (non-induced Vector Only control). We attributed this difference to the inhibitory effect of IPTG on cell viability (22). Additionally, this may reflect the fact that MTA is a metallothionein comprised of a large number of cysteine residues. Cysteines are the second most expensive amino acids to produce in *E. coli* based on the number of ATP molecules consumed to produce one molecule of it (23).

While the induced Vector Only controls and non-induced Ni_v.1 samples had a 14.3 ± 7.9% and 7.7 ± 1.7% reduction in their NGR, respectively, induced Ni_v.1 unexpectedly experienced a 45 ± 15.6% NGR reduction (Fig. 4D). Additionally, Ni_v.1 reproduced the same cell pellet appearance at the end of T5 as Fig.S3 in Test 1. We interpreted this difference as a consequence of burden-induced sensitization of the Ni_v.1’s cell membrane. Specifically, the T7-controlled overexpression of NikABCDE likely overwhelmed membrane protein biogenesis, disrupted homeostasis of the membrane’s fatty acid composition, and deteriorated the membrane integrity by physically over-occupying the membrane “real estate” (24–26). Studies have also showed that bacterial membranes and *E. coli* cells treated with EDTA led to significant outer membranes deformations (27, 28). Taken together, the behavior of Ni_v.1’s growth kinetics pointed to a need for the IPTG concentration to be re-calibrated for improved cell viability.

We confirmed IPTG, and not the nickel solution, imposed the larger burden based on growth curves of the Vector Only control and Ni_v.1 samples in the presence of only 10 ppm NiCl_2_, 0.4 mM IPTG, or both (Fig. S6). We observed the non-induced Vector Only controls and Ni_v.1 samples in the absence of nickel or IPTG had the fastest growth rates (Fig. S6A,B). The addition of 10 ppm NiCl_2_ did not lead to a significant change in growth rates. However, 0.4 mM IPTG delayed both the Vector Only control and Ni_v.1 samples’ exponential phase by ∼8-9 hrs. Ni_v.1 was slightly more delayed than the Vector Only control in the presence of nickel (Fig. S6C), but were both equally delayed in the presence of IPTG (Fig. S6D). Next, we tested the effect of different IPTG concentrations on the growth rates and protein expression levels to refine the workflow (Fig.2) for better Ni_v.1 viability.

### Test 3: Determining minimum IPTG concentrations required to induce NikA expression in Ni_v.1

Due to IPTG’s inhibitory effect on the Vector Only control and Ni_v.1 samples’ growth rates, we assessed the protein expression levels of the transport and storage system at various IPTG concentrations to determine its minimum order of magnitude required to achieve expression without major losses in cell viability. We obtained their growth curves at 0, 0.001, 0.01, 0.1, and 1 mM IPTG over a 16-hr period, and obtained 1 mL samples of the cultures every 4 hrs to assess the protein expression by SDS-PAGE analysis. We specifically tracked NikA (∼58 kDa) as a proxy for the expression of the transport and storage system. The ORF of *nikABCDE* was amplified for cloning using primers complementary to the NikA start codon (front) and the NikE stop codon (end), meaning *nikABCDE*’s native RBS for NikA was not kept when the full ORF was cloned into pCDF-Duet. Consequently, the NikA ORF in pCDF-Duet used the vector’s stronger RBS (based on the RBS calculator), and therefore had visibly higher expression than the NikBCDE permease complex (29). To compare samples at different IPTG concentrations and different timepoints more easily, we normalized the amount of protein loaded into each well to ensure the intensity of the positive control bands at ∼75 kDa were as consistent as possible on the SDS gel. In earlier purification work (unpublished), we found that NikA and its homologues would dimerize if they were re-solubilized from insoluble pellet fractions due to sub-optimal expression conditions, or if they were exposed to harsh conditions (*e*.*g*., high urea or guanidine concentrations). We therefore monitored bands at 116 kDa as well to assess NikA dimerization.

At 0 mM IPTG, Vector Only controls and Ni_v.1 samples predictably had the highest growth rates (Fig. 5A,B). Vector Only control growth rates were affected at 1 mM IPTG, whereas Ni_v.1 growth rates were slightly delayed by 0.1 mM IPTG and largely delayed by 1 mM IPTG. In the SDS-PAGE analysis of the Ni_v.1 cell lysates, the intensities of the bands at 75 kDa were consistent across the IPTG concentrations and timepoints, except for the 4hr and 8hr timepoints for 1 mM IPTG that should be interpreted more judiciously (Fig. 5C). At 0 mM IPTG, we observed no noticeable overexpression of NikA across the 16-hr period. Strong NikA overexpression appeared at 0.001 mM and 0.01 mM IPTG, with no dimerization observed at 116 kDa. Combined with the growth rate data, this suggests that a two-order magnitude of decrease in the original 0.4 mM IPTG concentration used in Test 1 and 2 may have been enough to express Ni_v.1’s transport and storage system without major losses in the cell viability. At 0.1 mM IPTG, NikA expression appears earlier at the 4hr timepoint compared to the lower IPTG concentrations, however at the 16hr timepoint NikA expression was weaker and its dimerization stronger. Given the Ni_v.1 growth rate (Fig. 5B) was reduced at 0.1 mM IPTG, the NikA dimerization likely reflected the increased burden. At 1 mM IPTG, NikA dimerization was more apparent with weaker NikA expression. Taken together, the IPTG concentrations used in previous studies was too high for Ni_v.1. Low micromolar levels of IPTG were sufficient for expression of NikA, and presumably the expression of the transport and storage systems.

**Figure 5.**
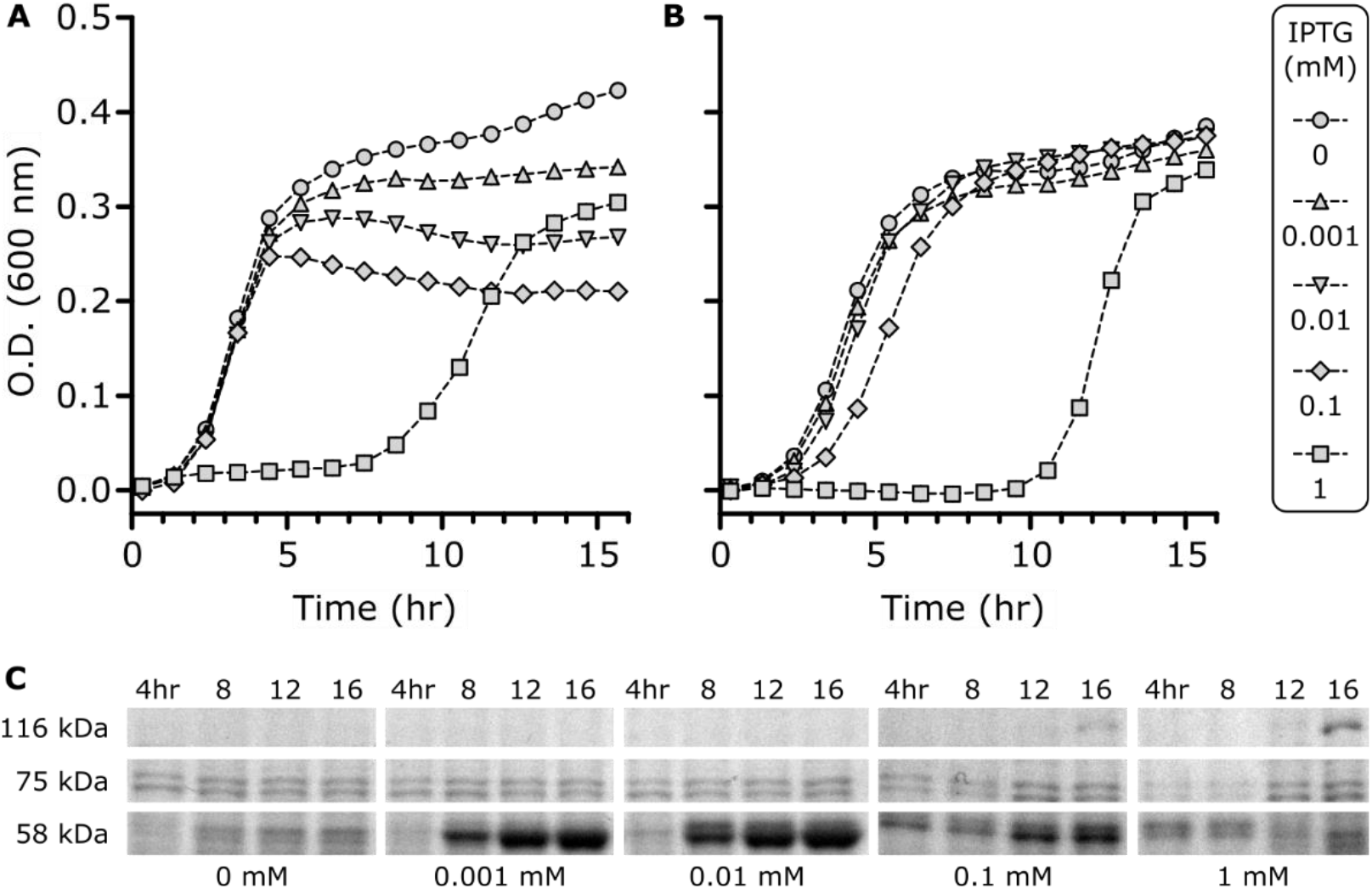
Ni_v.1 viability and protein expression as a function of IPTG concentration. (A) Growth curves of Vector Only controls and (B) Ni_v.1 samples (n=3, each). Every third datapoint was plotted to reduce clutter. Dotted lines are based on all data points. (C) SDS-PAGE analysis of Ni_v.1 cell lysates to monitor NikA expression. Samples are normalized based on their O.D. at each timepoint to ensure intensities of 75 kDa bands are consistent across samples in the SDS gel.

### Conclusion

In this study, we engineered an *E. coli* strain to have enhanced bioaccumulation performance for nickel. We first characterized aqueous streams originating from the Vale Copper Cliff site in Sudbury, Ontario to determine real-world constraints that Ni_v.1 would need to operate under. Then, through a series of tests, we demonstrated Ni_v.1 could remove nickel from a 10 ppm NiCl_2_ solution (matching the concentration found at the C.C. site) and achieved a bioaccumulation performance of 6.53 ± 1.47 mg_Ni_/g_dcw_, which was comparable to values published in the literature. In an analysis of the Vector Only control and Ni_v.1 samples’ growth kinetics, we found IPTG was likely causing burden-induced membrane sensitization in Ni_v.1 cells, and therefore increased cell lysis throughout the workflow outlined in Fig.2. Using a gradient of IPTG concentrations between 0-1 mM IPTG, we found low micromolar levels of IPTG were sufficient for expression of the transport and storage system needed to perform the nickel bioaccumulation task, despite previous studies using 0.4 - 1 mM IPTG.

In immediately subsequent DBTL cycles, Ni_v.1’s bioaccumulation performance for nickel in more complex solutions matching the working constraints of the N.C. and C.C. sites by first isolating for the effects of pH, salinity, and interfering metals such as Fe^2+/3+^, Co^2+^, and Cu^2+^ is needed. More rigorous experiments based on factorial designs should be used to systematically understand Ni_v.1’s performance in multi-parameter scenarios, which constitutes a novel approach to characterizing engineered strains in this field. Critically, the workflow outlined in Fig.2 needs to be miniaturized to facilitate more rapid DBTL cycles. Test 2 required 15 cultures that took a single person 12 consecutive hours to grow and fractionate, with an additional two day to process. In other work, we developed protocols on the Tecan Freedom Evo 100 to grow and analyze cell cultures in microplates that could be deployed in this work to automate T1→T3 and simplify the fractionation steps of T3→T6 (30).

In future DBTL cycles, additional objectives could focus on improving the growth kinetics. For example, plasmid burden could be alleviated by genomic integration of the NikABCDE transporter and MTA storage system. Further, functional stability could be improved by controlling a genome-integrated transport and storage system with a strong endogenous promoter to eliminate the need for inducing agents and reduce the metabolic burden associated with T7-based expression. Another objective could be to develop a new genetic system that enables the cell to export the nickel once it has reached a “saturation threshold” such that the cell can be re-used for multiple metal recovery cycles, thus improving its economics. Finally, this was the first demonstration of a nickel-specific ABC transporters heterologously expressed for the purpose of removing nickel from solution. Protein engineering efforts could be undertaken to explore the possibility of altering NikA’s metal specificity – and therefore the strain’s specificity – for other metals with strategic value (*e*.*g*., precious metals and rare earth elements) towards stronger supply chains for advanced clean technologies and circular economies.

## MATERIALS AND METHODS

### Aqueous stream sample collection and compositional analysis

The sampling sites were at Nolin Creek (46°28’21.9”N 81°03’53.9”W; 46.472759, -81.064958) and a Copper Cliff site (46°28’21.9”N 81°03’53.9”W; 46.472759, -81.064958) in Sudbury, Ontario, Canada. Eight 16 oz Nalgene bottles were acid washed in a 10% hydrochloric acid solution, rinsed with deionized water, dried, and weighed. Before taking samples from the two sites, litmus paper was used to determine the pH of the locations. The bottles were filled with water samples on July 28^th^, 2021, where one of each set was acidified with 25 drops of trace metal grade nitric acid for ICP analysis. Samples were packed in a cooler, with freezer packs, and shipped to University of Toronto on the same day. The water samples collected were stored in a walk-in fridge at 4 °C. To prepare for the elemental analysis, the samples were vacuum filtered (50µm pore size) to remove solid sediments, and then diluted (5 folds) in 5% HNO_3_ to stabilize the content. The sample pH values were measured before and after the vacuum filtration, which remained largely constant. Common cation elements (Al, K, Pb, Mn, Cd, Fe, Mg, Na, Co, Zn, Cu, Cr, Ca, and Ni) (Inorganic Ventures, #IV-28) and suspected anion elements (S, P, and As) (Inorganic Ventures, #CGS1; High-Purity Standards, #100039-1; Inorganic Ventures, #IV-28) were measured via ICP-OES. A chloride probe was used to determine the Cl^-^ concentration. Because the water samples were collected at the ground level and have been exposed to atmosphere extensively, it is assumed that all Fe exists as ferric ion, and all S exists as sulfate ion in the valency calculations and subsequent charge balances.

### Plasmid preparation and cloning workflow

The open-reading frame of the *nikABCDE* gene was amplified from genomic DNA belonging to *Escherichia coli* BL21(DE3) using the KAPA-HiFi Master Mix kit according to the manufacturer’s instructions (Kapa Biosystems, #KK2601). Primers are found in Supplementary Table 3. The forward primer (#0) included the NcoI cut site, and the reverse primer (#1) included the HindIII cute site. These cut sites were chosen to remove the 6x-HisTag and the majority of the other cut sites. The open-reading frame of the *mta* gene was synthesized through Twist Biosciences (San Francisco, United States) and amplified in the same fashion. Specifically, MTA’s forward primer (#2) included an NdeI cut cite and the reverse primer (#3) a PacI cut site for the same reasons as primers #1 & 2. All restriction enzymes were obtained from New England Biolabs. Amplified products were gel-extracted and cleaned using a column prep kit (ThermoFisher, # K0701). Restriction digest procedures followed the manufacture protocol. The dual expression vector pCDF-Duet (Novagen: 71340) was linearized using the appropriate restriction enzyme pair for NikABCDE or MTA, gel-extracted, and cleaned using the column prep kit. T4 DNA ligase (NEB, # M0202) was used to ligate NikABCDE or MTA into the appropriate linearized pCDF-Duet vector, then transformed into in-house calcium-competent *E. coli* strains DH5α and LOBSTR-BL21(DE3) (Kerafast, #EC1002). NikABCDE was cloned into the first site, and MTA into the second site. To create the full transport-storage system (Ni_v.1), MTA was cloned into the second site of the vector already containing NikABCDE in the first site. The completed plasmids were miniprepped (ThermoFisher, #K0502) and sequence verified at ACGT Corp (Toronto, Canada) using primers #4-7. Glycerol stocks were stored at - 80°C. Links for the final constructs are found in Supplementary Table 4.

### Shake flask-based determination of bioaccumulation performance with ICP-OES

Starter cultures of *E*.*coli* Ni_v.1 were grown overnight in 10 mL of sterile Luria-Bertani media (BioShop: LBL407.1) containing 100 μg/mL of streptomycin for 16 hr at 37 °C with shaking (200 rpm). Bioaccumulation cultures were prepared by inoculating 100 mL of sterile Luria-Bertani media containing 100 μg/mL streptomycin with a 1% v/v inoculum of the starter culture and growing for 3 hrs at 37 °C with shaking. Isopropyl β-d-1-thiogalactopyranoside (IPTG) was added to a final concentration of 0.4 mM to induce the expression of the NikABCDE and MTA proteins. Bioaccumulation cultures were grown for another 3 hrs at 37 °C with shaking, then centrifuged (5000 rpm, 10 min, 25 °C) into conical vials (50 mL). The supernatant (*i*.*e*., spent media) was discarded and the cell pellet was re-suspended in 10 mM HEPES pH 7.2 to rinse the media. The cells were centrifuged again and the supernatant (*i*.*e*., HEPES buffer) was discarded. The cells were then resuspended in 26.4 mL of 10 ppm NiCl_2_ prepared in MilliQ water and allowed to bioaccumulate Ni^2+^ for 3 hrs at 37 °C with shaking. Afterwards, the mixtures were individually transferred into smaller conical vials (15 mL) and centrifuged to collect the biomass. This was repeated for the remaining mixture. The supernatants (residual fraction) were collected in new 50 mL conical vials and stored at 4 °C. If a washing step was completed, the cells were then re-suspended in 5 mL 10 mM EDTA, then centrifuged again where the supernatant (wash fraction) was stored at 4 °C in new vials. Whether a wash step was completed or not, the smaller conical vials were pre-weighed to determine their mass of the pellet. Finally, the cell pellet (washed or not washed) was freeze-dried for 24 hr, then re-suspended in 5 mL 70% HNO_3_^-^ (FisherScientific: A509P500) for a 24 hr acid digestion period (biomass fraction). Fractions (*i*.*e*., residual, wash, and biomass) were analyzed by inductively coupled plasma optical emission spectrometry (ICP-OES, Agilent Dual View 720) using a Ni^2+^ calibration curve with 0, 1, 5, 10, and 20 ppm standards.

### Microplate-based determination of cellular growth rates

Growth curves were obtained by tracking the optical density of cells as they grew in 24-well microplates using the Infinite® M200 (Tecan) plate reader. All sample were grown at 37 °C with shaking (orbital, 2mm) for 16 hrs with 20-min interval reads. To prepare each well for inoculation, sterile room temperature LB was first pre-mixed with streptomycin to 100 μg/mL. IPTG (0.4 mM for Test 2 [Fig. S4], 0→1 mM for Test 3 [Fig. 6]) and NiCl_2_ (10 ppm for Test 2 [Fig. S4]) were added as needed to achieve conditions necessary for experiments. 1000 μL of the mixtures were aliquoted into each well. 5 μL of overnight culture (grown to saturation) were inoculated into the wells as prescribed by the experiment design.

### SDS-PAGE analysis of Ni_v.1 cell lysates and NikA overexpression

In parallel with cells grown in microplates to obtain their growth curves (Fig. 6), 10 mL LB cultures (pre-mixed with the necessary components as described above) of the same cells were grown in glass test tubes. 1 mL aliquots were taken from cultures every 4 hrs for a 16hr period. The optical densities for each set of aliquots at each time point were measured in cuvettes, then centrifuged to collect the cell pellet (supernatant discarded). Cell pellets were frozen at -20 °C until all pellets were collected after the full 16hr period. For a fair comparison of NikA expression (∼58 kDa) across the different IPTG concentrations and timepoints, we normalized the dilution of the cell pellets in BugBuster lysis buffer (Millipore Sigma, #70584) such that the approximate concentration of cells was the same as the 1 mM IPTG induced Ni_v.1 sample. We then created mixtures comprised of 20 μL cells + 5 uL of 6X SDS dye, boiled the mixtures for 10 min, and loaded 15 μL of the mixtures into wells.

## ACKNOWLEDGEMENTS

The authors thank Elizabeth Edwards for her helpful feedback on experiments and data interpretations, as well as Indje Mihaylov and Glen Watson from Vale for their useful input. This work was supported by the Ontario Ministry of Economic Development, Job Creation and Trade through the Elements of Bio-mining ORF-RE program. PD is grateful to be a recipient of Ontario Graduate Scholarships. PD: conceptualization, methodology, investigation, formal analysis, original draft, editing. HS, JAW: investigation, formal analysis, original draft. NM, AFY: supervision, editing. VP, RM: supervision, review, editing.

## ABBREVIATIONS

ABC: ATP-binding cassette (transporter)
SBP: solute-binding protein
IPTG: isopropyl ß-D-1-thiogalactopyranoside
NTP: nucleotide triphosphate
ICP-OES: inductively coupled plasma optical emission spectrometry
LB: Luria-Bertani (nutrient media)
kDa: kilodaltons
O.D.: optical density
NGR: normalized growth ratio

