## Supplementary Materials for "Engineered nickel bioaccumulation in *Escherichia coli* by NikABCDE transporter and metallothionein overexpression"

Supplementary Table 1. **Summary of the elements from the two sampling sites based on ICP-OES analysis (n=3)**

| Elements | Units | Al | K | Pb | Mn | Cd | Fe |
|---|---|---|---|---|---|---|---|
| N. C. | mmol/L | 0.004 ± 0.004 | 0.276 ± 0.009 | n.a. | 0.001 ± 0.0003 | n.a. | 0.011 ± 0.001 |
| | mg/L | 0.106 ± 0.110 | 10.790 ± 0.354 | n.a. | 0.051 ± 1.564 | n.a. | 0.597 ± 0.054 |
| C. C. | mmol/L | 0.190 ± 0.024 | 0.824 ± 0.074 | n.a. | 0.028 ± 0.003 | n.a. | 1.163 ± 0.139 |
| | mg/L | 4.378 ± 0.655 | 32.211 ± 2.922 | n.a. | 1.564 ± 0.167 | n.a. | 64.937 ± 7.761 |
| | | **Mg** | **Na** | **Co** | **Zn** | **Cu** | **Cr** |
| N. C. | mmol/L | 0.161 ± 0.018 | 0.452 ± 0.028 | n.a. | n.a. | 0.003 ± 0.001 | n.a. |
| | mg/L | 3.913 ± 0.433 | 10.38 ± 0.655 | n.a. | n.a. | 0.162 ± 0.089 | n.a. |
| C. C. | mmol/L | 2.540 ± 0.254 | 2.560 ± 0.261 | 0.006 ± 0.001 | 0.005 ± 0.0004 | 0.016 ± 0.001 | n.a. |
| | mg/L | 61.725 ± 6.128 | 58.848 ± 6.005 | 0.380 ± 0.064 | 0.301 ± 0.029 | 0.998 ± 0.092 | n.a. |
| | | **Ca** | **Ni** | **S** | **P** | **As** | **Cl** |
| N. C. | mmol/L | 0.258 ± 0.038 | 0.005 ± 0.001 | 0.848 ± 0.022 | 0.027 ± 0.0001 | n.a. | 0.001 ± 0.000 |
| | mg/L | 10.321 ± 1.518 | 0.268 ± 0.045 | n.c. | n.c. | n.a. | n.c. |
| C. C. | mmol/L | 6.393 ± 0.878 | 0.184 ± 0.019 | 13.856 ± 1.53 | 0.026 ± 0.001 | n.a. | 0.002 ± 0.000 |
| | mg/L | 256.235 ± 35.196 | 10.805 ± 1.118 | n.c. | n.c. | n.a. | n.c. |

N.C. – Nolin Creek; C.C. - Copper Cliff.
n.a. entry represents data below detection limit or below 1 µmol/L; n.c. entry was not calculated.

Supplementary Table 2. **Nickel fractionation data (correspoding to Test 2)**

| Fraction | pCDF-Duet (Empty Vector) | | | | Ni_v.1 | | | |
|---|---|---|---|---|---|---|---|---|
| | +IPTG | | -IPTG | | +IPTG | | -IPTG | |
| | % Ni | $mg_{Ni}$ | % Ni | $mg_{Ni}$ | % Ni | $mg_{Ni}$ | % Ni | $mg_{Ni}$ |
| Biomass [a] | 13.5 ± 1.5 | 0.015 ± 0.001 | 29.9 ± 2.2 | 0.032 ± 0.002 | 23.2 ± 1.6 | 0.025 ± 0.002 | 28.0 ± 3.2 | 0.030 ± 0.004 |
| Wash [b] | 35.6 ± 0.4 | 0.040 ± 0.001 | 32.0 ± 1.9 | 0.035 ± 0.002 | 23.0 ± 0.8 | 0.025 ± 0.002 | 31.8 ± 2.9 | 0.034 ± 0.003 |
| Residual | 50.9 ± 1.5 | 0.057 ± 0.002 | 38.1 ± 1.7 | 0.041 ± 0.002 | 53.7 ± 2.2 | 0.058 ± 0.001 | 40.2 ± 9.3 | 0.043 ± 0.001 |

Percentage nickel found in each fraction was calculated by dividing the $mg_{Ni}$ in each fraction by the sum of the $mg_{Ni}$ measured across all fractions for each strain (empty vs. Ni_v.1) and condition (IPTG).
[a] – the amount of nickel bioaccumulated into the cell; [b] – the amount of nickel biosorped on the surface of the cell

Supplementary Table 3. **Primers**

| ID | Primer Name | Sequence (5’ to 3’) | $T_m$ (°C) | Purpose |
|---|---|---|---|---|
| 0 | Forward NikABCDE | gtacgaccatggatgctctccacactccgccgcactc | 65 | Cloning |
| 1 | Reverse NikABCDE | gtacgaaagcttttaaaccttttctgtggtgcgacggcgca | 65 | |
| 2 | Forward MTA | gtacgacatatgagcggctgcggttgcggcag | 65 | Cloning |
| 3 | Reverse MTA | gtacgattaattaattacttacagttgcagggatcacaggtgcaattatcaccgc | 65 | |
| 4 | Forward Site 1 Sequencing | ggatctcgacgctctccct | 57 | Sequencing |
| 5 | Reverse Site 1 Sequencing | gattatgcggccgtgtacaa | 55 | |
| 6 | Forward Site 2 Sequencing | ttgtacacggccgcataatc | 55 | Sequencing |
| 7 | Reverse Site 2 Sequencing | gctagttattgctcagcgg | 53 | |

Supplementary Table 4. **Sequences**

| Sequence Name | Link to Sequences [b] |
|---|---|
| pCDF-Duet Vector | https://benchling.com/s/seq-PblPbPGll2SVQzCKoDEF?m=slm-SB4QqukyRhd6Wndg6xTJ |
| MTA Only | https://benchling.com/s/seq-UbCXMDVrU3IJEIYdKr76?m=slm-0qaIb0Xpj0AbycIyurIc |
| NiKABCDE Only [a] | https://benchling.com/s/seq-mWaP9Yy6MfSXuzQu8F7k?m=slm-0oHRljWjtp56UXqM0cjd |
| Full Ni_v.1 Construct [a] | https://benchling.com/s/seq-JyRqP9nA1wXe4l7plAcq?m=slm-SIvaxbfhQWLuo9YHK09f |

[a] An additional base pair leading to a frameshift mutation for NikABCDE and the full Ni_v.1 construct was corrected (*i.e.*, removed) by Ranomics through site directed mutagenesis prior to all testing.

[b] If the URL links do not work, please contact the corresponding author.

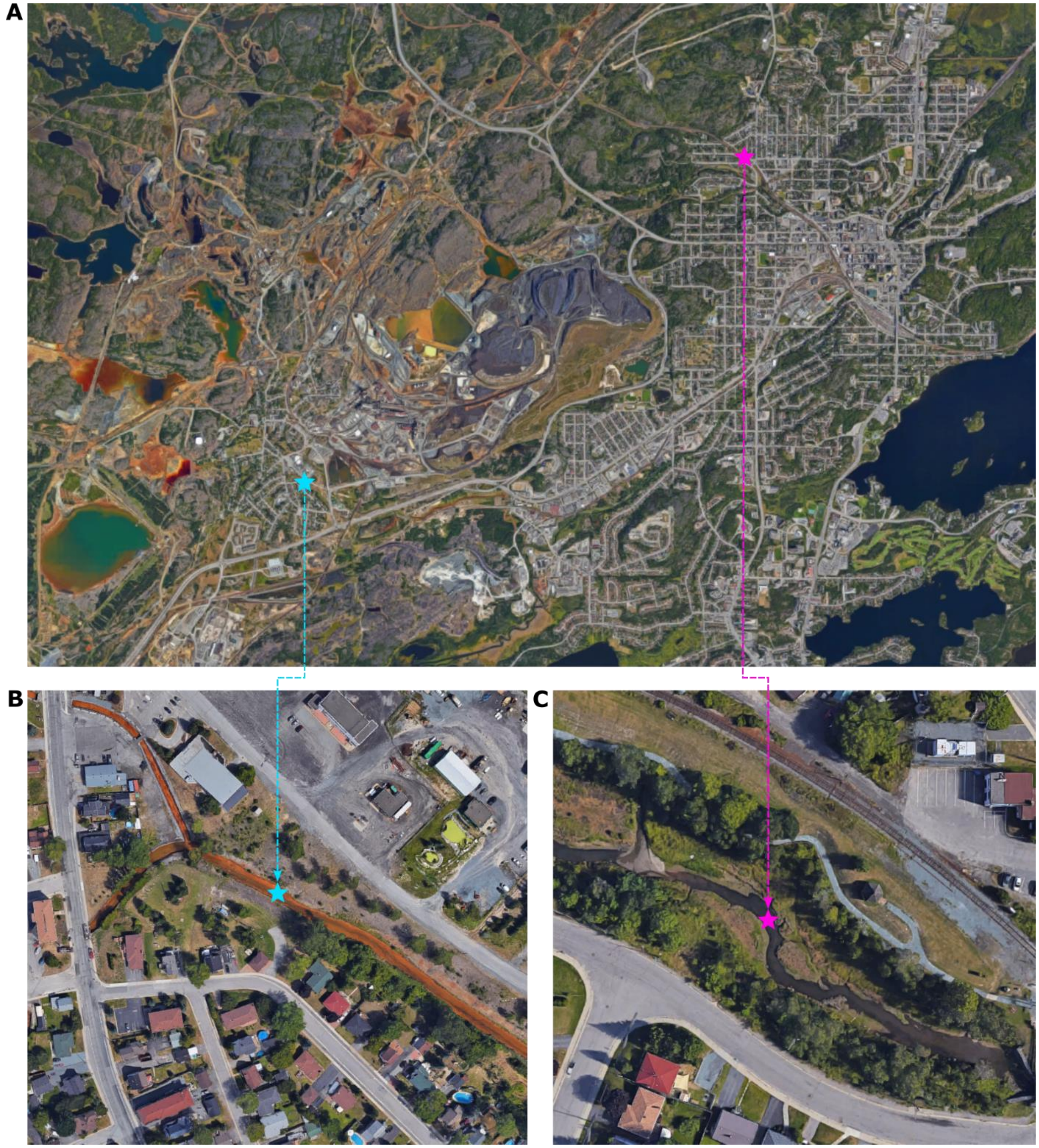


Supplementary Figure 1. **Aerial view of aqueous stream sampling sites**. Blue star – Copper Cliff (C.C.). Pink star – Nolin Creek (N.C.). (A) View of Vale Copper Cliff site and the City of Greater Sudbury. (B) View of the C.C. site. (C) View of the N.C. site. Aerial images taken from Google Earth (accessed on July 2022).

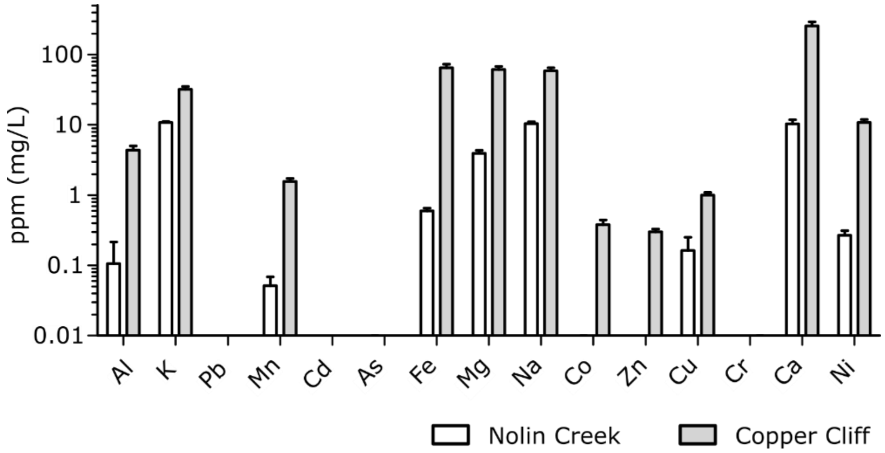


Supplementary Figure 2. **Metal content comparison between sampling sites.** Same data displayed in Figure 3 of main text, but with metal concentrations for each metal plotted side-by-side for easier comparisons.

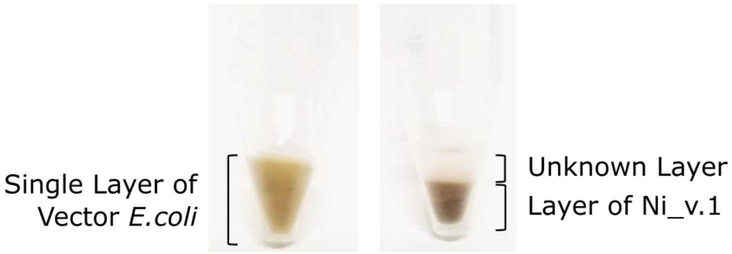


Supplementary Figure 3. **Comparison of pellet appearance before acid digestion at the end of T5.** Vector Only control (left) and Ni_v.1 (right) pelleted via centrifugation in 15 mL conical vials.

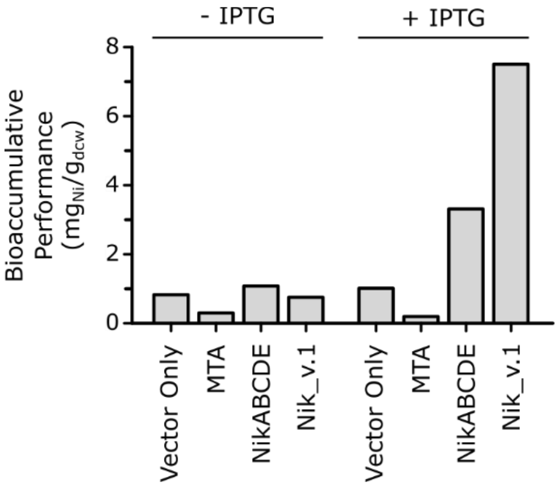


Supplementary Figure 4. **Ni_v.1 bioaccumulation performance compared to controls.** Single replicate flask experiments for four strains were tested in the absence and presence of 0.4 mM IPTG for induction: Vector Only (empty pCDF-Duet vector where both storage and transport systems were absent), MTA (only the storage system was present), NikABCDE (only the transport system was present), and Nik_v.1 (where both the transport and storage systems were present).

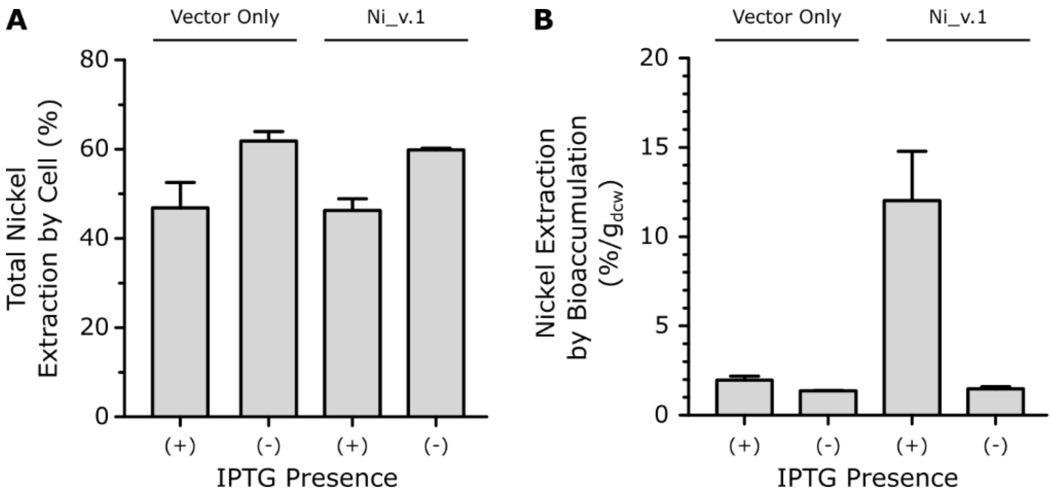


Supplementary Figure 5. **Nickel extraction performance.** Vector Only, referring to the strain carrying the empty pCDF-Duet vector, and Ni_v.1 referring to the strain carrying both NikABCDE and MTA. (A) Percent nickel extraction based on the total nickel ($mg_{Ni}$) measured across all three fractions (residual, wash, and biomass). (B)

Percent nickel extractions normalized to the amount of cells ($g_{DCW}$) obtained in the biomass fraction. Experiments were performed in triplicate (n=3).

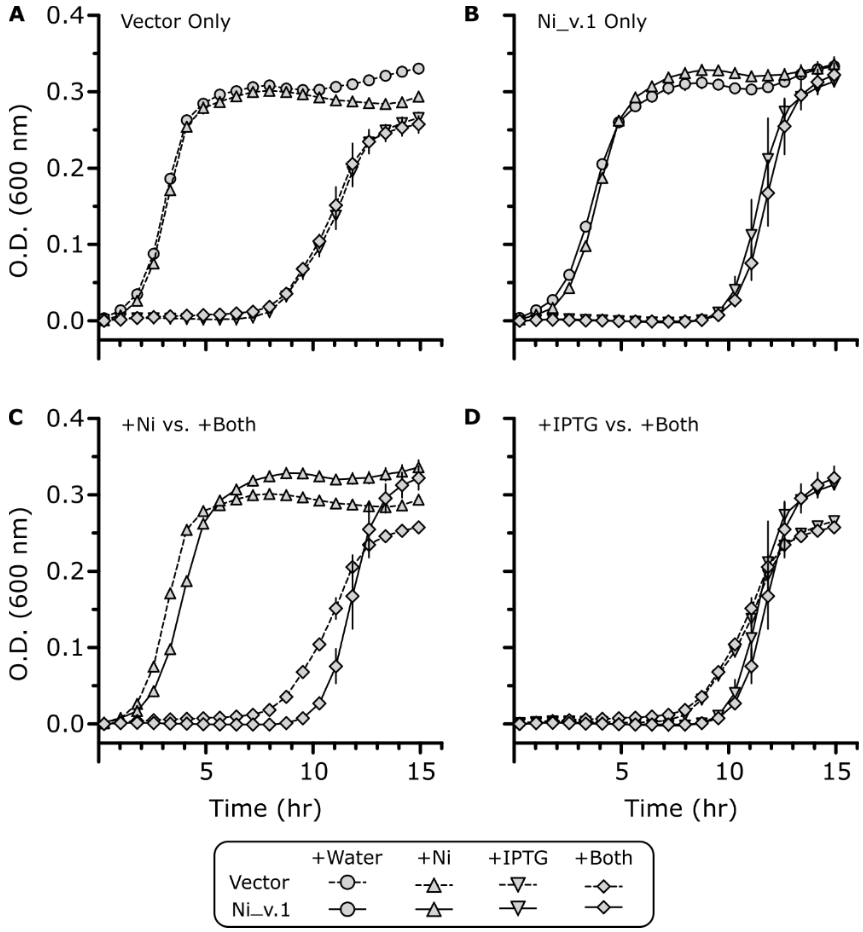


Supplementary Figure 6. **Assessment of Ni_v.1 growth inhibition by nickel and IPTG**. Growth curves are plotted to compare the effects of 10 ppm $NiCl_2$, 0.4 mM IPTG, or both for (A) Vector Only controls, and (B) Ni_v.1 samples. The synergistic effects of nickel (C) and IPTG (D) are compared (n=3). Every other three datapoints were plotted to avoid cluttering.
